# Distinct members of the *C. elegans* CeMbio reference microbiota exert cryptic virulence and infection protection

**DOI:** 10.1101/2023.11.02.565327

**Authors:** Xavier Gonzalez, Javier E. Irazoqui

**Affiliations:** Immunology and Microbiology graduate program, Morningside Graduate School of Biomedical Sciences, University of Massachusetts Chan Medical School, Worcester MA 01605; Department of Microbiology and Physiological Systems, University of Massachusetts Chan Medical School, Worcester MA 01605

## Abstract

Microbiotas are complex microbial communities that colonize specific niches in the host and provide essential organismal functions that are important in health and disease. A key aspect is the ability of each distinct community member to promote or impair host health, alone or in the context of the community, in hosts with varied levels of immune competence. Understanding such interactions is limited by the complexity and experimental accessibility of current systems and models. Recently, a reference twelve-member microbiota for the model organism *C. elegans,* known as CeMbio, was defined to aid the dissection of conserved host-microbiota interactions. Understanding the physiological impact of the CeMbio bacteria on *C. elegans* is in its infancy. Here, we show the differential ability of each CeMbio bacterial species to activate innate immunity through the conserved PMK-1/p38 MAPK, ACh/WNT, and HLH-30/TFEB pathways. Using immunodeficient animals, we uncovered several examples of bacterial ‘cryptic’ virulence, or virulence that was masked by the host defense response. The ability to activate the PMK-1/p38 pathway did not correlate with bacterial virulence in wild type or immunodeficient animals. In contrast, ten out of twelve species activated HLH-30/TFEB, and most showed virulence towards *hlh-30-*deficient animals. In addition, we identified *Pseudomonas lurida* as a pathogen in wild type animals, and *Acinetobacter guillouiae* as avirulent despite activating all three pathways. Moreover, short pre-exposure to *A. guillouiae* promoted host survival of infection with *P. lurida,* which was dependent on PMK-1/p38 MAPK and HLH-30/TFEB. These results suggest that the microbiota of *C. elegans* is rife with “opportunistic” pathogens, and that HLH-30/TFEB is a fundamental and key host protective factor. Furthermore, they support the idea that bacteria like *A. guillouiae* evolved the ability to induce host innate immunity to improve host fitness when confronted with pathogens, providing new insights into how colonization order impacts host health.

## Introduction

Animals exist as super-organisms, in association with complex and dynamic microbial communities known as the microbiota (Eberl, 2010; Kostic *et al*., 2013; Peixoto *et al*., 2020). Although awareness of the importance of animal microbiotas emerged decades ago (Marples *et al*., 1970; Marples and Kligman, 1971; Shilov *et al*., 1971), the local and systemic influences of the microbiota on host physiology have come into focus only in recent times (Huttenhower *et al*., 2012; Peixoto *et al*., 2020). Although examples of specific mechanisms of microbiota-host interaction have emerged (Kamada *et al*., 2013; Gasaly *et al*., 2021; Caballero-Flores *et al*., 2023; Horrocks *et al*., 2023), many aspects remain poorly understood. In particular, the functional significance of distinct microbiota members, in terms of their microbe-microbe and microbe-host interactions, is not fully understood. Due to the sheer size of such communities (in the order of thousands of species in humans (Wang *et al*., 2017)) and to a dearth of information regarding the microenvironments within the host in which these communities assemble and interact, it is extremely challenging to develop a comprehensive understanding of microbe-microbe and microbiota-host interactions in health and disease. Smaller reference communities (Lawley *et al*., 2012; Kostic *et al*., 2013; Brand *et al*., 2015) and simpler model organisms can be of assistance in understanding such fundamental principles and the evolutionary forces at play.

To fully understand the emerging properties of such complex microbial communities in interaction with the host requires moving beyond phenotypic potentials that are encoded in the individual assembled genomes and meta-genomes to actual phenotypes, the expression of which is highly context-dependent. To develop a better understanding of microbe-microbe and microbe-host interactions, one approach is to define the phenotypic states of each microbiota member in isolation, within various environments and including distinct host physiological states. Host mono-colonization is frequently used as an approach to unravel host-microbiota interactions, including interactions with the host’s immune system (Geva-Zatorsky *et al*., 2017; Rogala *et al*., 2020; Weitekamp *et al*., 2021). With current technologies, this problem quickly becomes intractable due to the vast array of community members, the diverse potentials their genomes encode, variable environments, and variation in host physiology across time and space.

*Caenorhabditis elegans* is an invertebrate model organism that associates with a much simpler intestinal microbiota than vertebrates, yet presents an innate immune system that is by and large conserved in higher organisms (Irazoqui *et al*., 2010b; Harding and Ewbank, 2021). Conserved signaling pathways, including the PMK-1/p38 MAPK, ACh-WNT, and HLH-30/TFEB pathways, connect the presence of pathogenic bacteria (or the damage they cause) with the induction of host defense genes that are also evolutionarily conserved (Irazoqui *et al*., 2010a; Visvikis *et al*., 2014; Labed *et al*., 2018; Fletcher *et al*., 2019). Although much has been accomplished to reveal and understand these pathways in the context of clinically relevant bacterial infections (e.g., *Pseudomonas aeruginosa, Enterococcus faecalis,* and *Staphylococcus aureus*) their roles in interactions with the microbes that *C. elegans* encounters in the wild and that constitute its natural microbiota have been largely unaddressed. The simplicity and ease of experimental manipulation of the model enables a reductionist approach to investigate microbe-microbe and microbe-host interactions in the whole, live organism.

Despite recent progress, much is unknown about the resident intestinal microbiota of *C. elegans,* in terms of its members, their interactions within the community, and their effects on host physiology and behavior. Recent groundbreaking studies have revealed mechanisms by which the microbiota affects host metabolism, gene expression, and behavior (Berg *et al*., 2016; Berg *et al*., 2019; Ortiz *et al*., 2021; Taylor and Vega, 2021; Obeng *et al*., 2023). However, which microbiota members are beneficial or harmful to the host under different environmental conditions remains poorly defined. To aid in answering such fundamental questions, a consortium of *C. elegans* researchers defined a minimal reference microbiota, named CeMbio, which comprises twelve representative species of bacteria from various clades that are frequently found in association with the wild *C. elegans* intestinal epithelium (Dirksen *et al*., 2020). How these microbiota members interact with the host’s innate immune system and how they affect each other is not well defined.

To address these knowledge gaps, we determined the ability of each CeMbio member to activate three important host defense pathways, namely the PMK-1/p38 MAPK, ACh-WNT, and HLH-30/TFEB pathways, and to cause disease in animals that lacked them. We identified a broad range of phenotypes both in terms of pathway reporter gene induction and virulence in wild type and immunodeficient animals. Furthermore, we identified a frank pseudomonad pathogen and an immunity-promoting Acinetobacter species, which demonstrated immune-mediated antagonism against the former. These microbe-microbe interactions suggest that the order of host colonization may be critical to host health. These seminal discoveries set the foundation for further in-depth investigation of the mechanisms that mediate “cryptic” virulence, or virulence that is masked by the host immune response, and “Third-Party” microbe-microbe interactions that are mediated through the host’s immune system.

## Experimental Results

### Growth characteristics of representative C. elegans microbiota species

We obtained single colonies of CeMbio bacteria on TSA medium and observed their macroscopic features (**Fig. 1A**). At 25 °C, most species formed colonies by 24 h, except *Pseudomonas berkeleyensis* and *Sphingomonas molluscorum,* which required up to 48 h. Most bacteria formed circular, smooth, opaque, and smooth-edged colonies, with the exception of *Comamonas piscis*, which formed undulated colonies. In most cases, colonies were shades of yellow-beige, except for *Chryseobacterium scophthalmum* (orange) and *Enterobacter hormaechei* (white) (**Fig. 1A**); after three weeks at 4 °C, *C. scophthalmum* colonies turned dark brown and secreted a dark pigment that diffused throughout the medium. Microscopically, the CeMbio bacteria were Gram-negative bacilli, except for *C. piscis,* which were cocci or coccobacilli (**Fig. 1B**).

**Figure 1.**
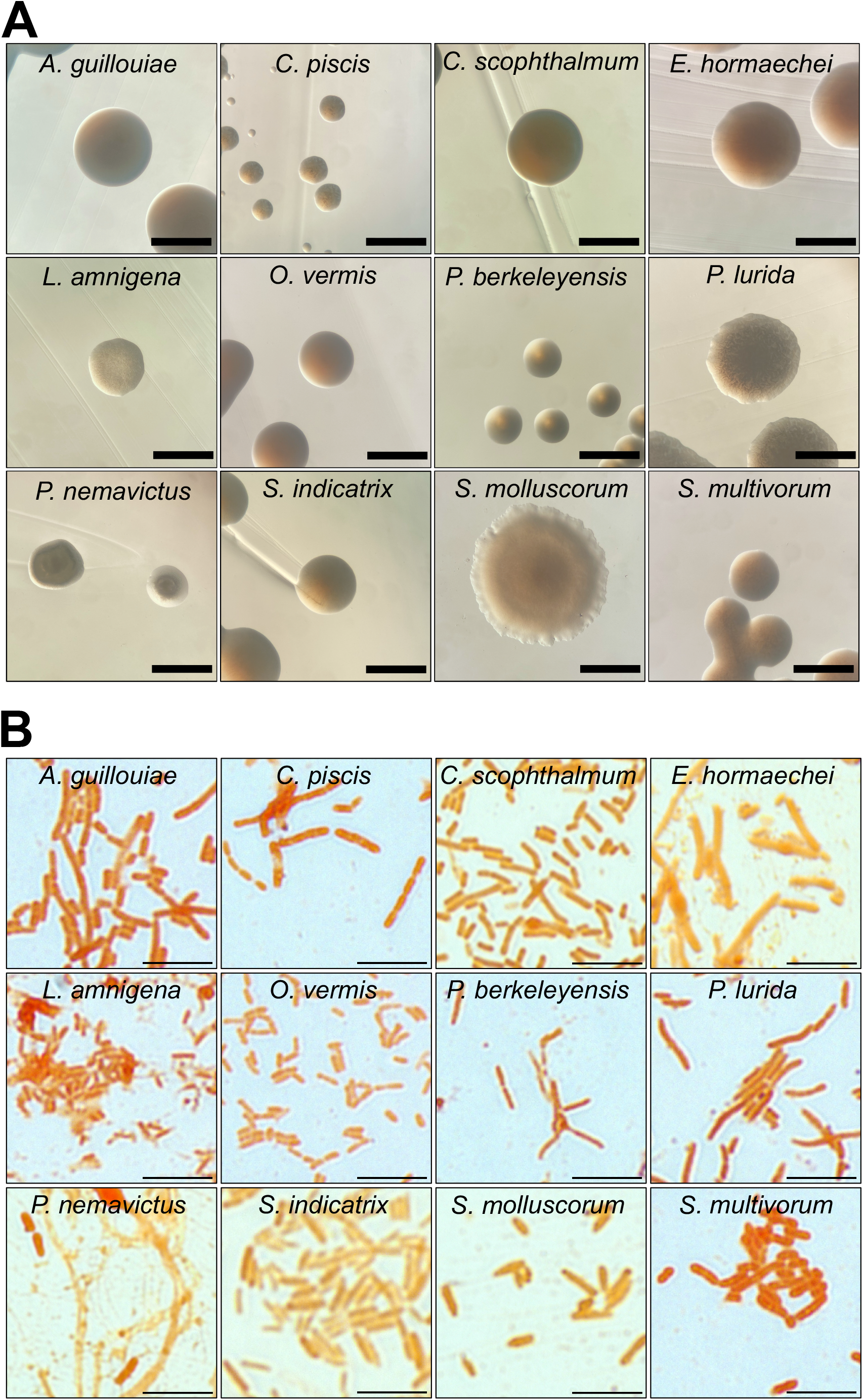
Growth and morphological characteristics of CeMbio bacteria. **A.** Representative images of individual CeMbio colonies grown on TSA at 25 °C for 24-48 h. Scale bars = 1 mm. **B.** Gram staining of individual CeMbio cultures grown overnight in TSB at 25 °C for 24 h. Scale bars = 5 μm.

The CeMbio species also varied in their antibiotic susceptibility. We tested two concentrations each of ampicillin, erythromycin, kanamycin, and streptomycin (**Fig. S1A and Table 1**). For ampicillin, all the CeMbio bacteria were resistant at 50 μg ml^-1^, and only *S. molluscorum* was susceptible to 50 μg ml^-1^. For erythromycin, all the species were resistant at 10 μg ml^-1^, except for *S. molluscorum* and *C. piscis.* At 50 μg ml^-1^, only *E. hormaechei*, *S. indicatrix*, *Lelliottia amnigena*, and *P. lurida* were also resistant. For kanamycin, *Sphingobacterium multivorum*, *Stenotrophomonas indicatrix*, *C. scophthalmum*, and *Ochrobactrum vermis* were resistant to 50 μg ml^-1^, while *P. lurida* was only resistant to 10 μg ml^-1^. For streptomycin, *S. multivorum*, *S. indicatrix*, *C. scophthalmum*, *S. molluscorum*, and *O. vermis* were resistant to 50 μg ml^-1^, while *C. piscis*, *P. berkeleyensis*, and *P. lurida* were resistant only to 10 μg ml^-1^. Thus, The CeMbio bacteria show extensive and partially overlapping antibiotic resistance. Three species (*O. vermis*, *S. multivorum,* and *C. scophthalmum*) exhibited resistance to all but 50 μg ml^-1^ erythromycin, and *S. indicatrix* was resistant to all. These results suggested that these *C. elegans-*associated bacteria are under strong selective pressure from antibiotics in the wild.

### CeMbio bacteria differentially activate a PMK-1/p38-dependent gene reporter in the intestinal epithelium

The p38 MAPK pathway is the best understood host defense pathway in *C. elegans* (Kim *et al*., 2002; Irazoqui *et al*., 2010b). To better delineate its role during interactions with individual members of the microbiota, we measured the expression of a genetically-encoded fluorescent construct that is used to monitor its activity (Shivers *et al*., 2010). This P*t24b8.5::gfp* reporter is induced in a *pmk-1-*dependent manner in the intestinal epithelium during intestinal infection by pathogens such as *P. aeruginosa* (Shivers *et al*., 2009). Reference animals fed nonpathogenic *E. coli* on NGM media showed P*t24b8.5::gfp* expression in the anterior and posterior ends of the intestinal epithelium, with little basal expression in between (**Fig. 2A, E**). Animals fed *E. coli* on TSA showed increased P*t24b8.5::gfp* expression along the entire intestinal epithelium (**Fig. 2E, H**). Animals on TSA media alone, or infected with *S. aureus* on the same, showed decreased P*t24b8.5::gfp* expression relative to reference animals, ruling out a nonspecific effect of the media and showing pathogen specificity of the reporter (**Fig. 2E, H**).

**Figure 2.**
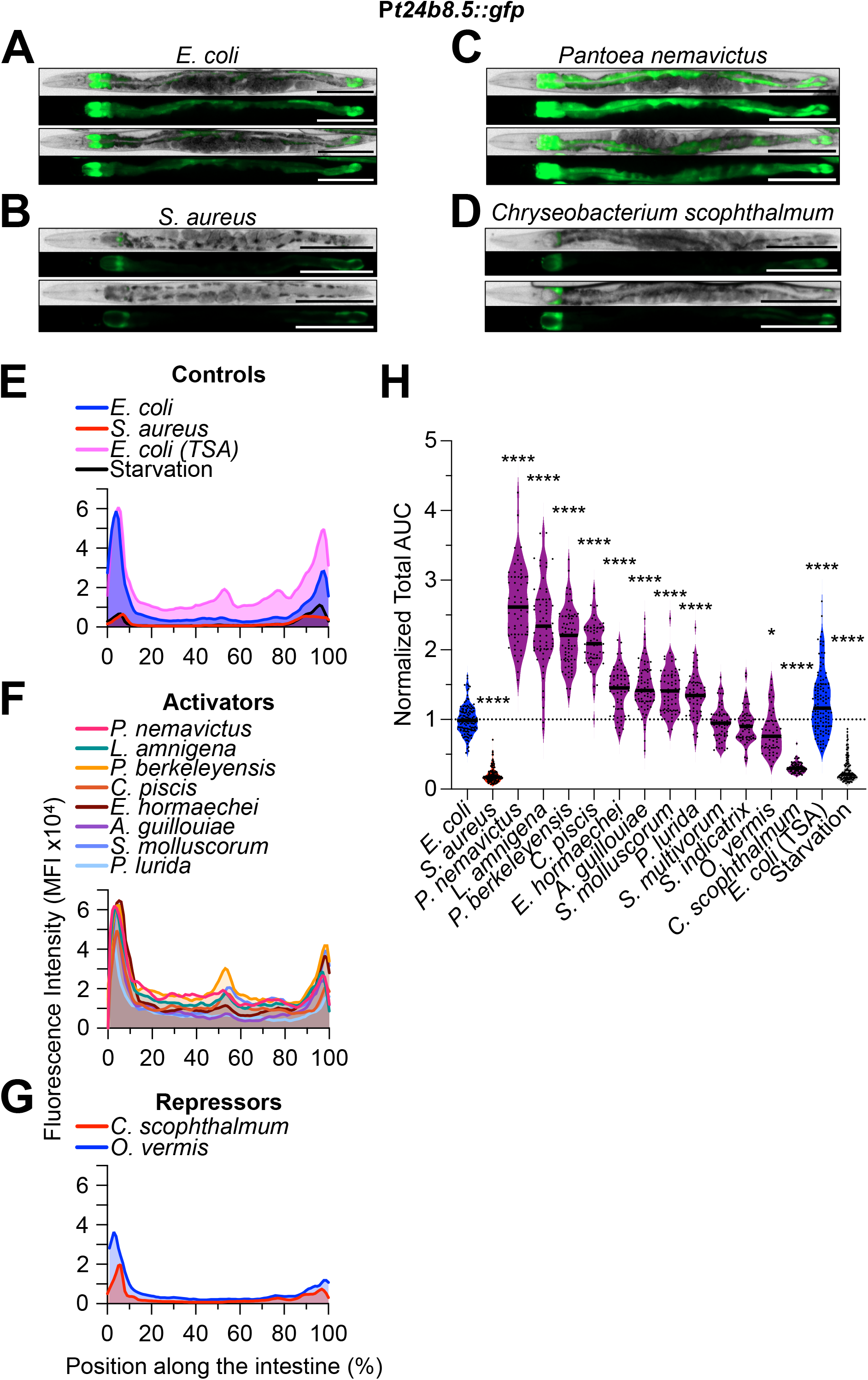
*C. elegans* microbiota bacteria differentially activate P*t24b8.5::gfp* expression along the intestinal epithelium. **A - D.** Representative brightfield and epifluorescence micrographs of P*t24b8.5::gfp* animals fed *E. coli*, *S. aureus*, *P. nemavictus*, and *C. scophthalmum* for 24 h at 25 °C. Animals were straightened using FIJI. Scale bars = 200 μm. **E - G.** P*t24b8.5::gfp* mean fluorescent intensity (MFI) along the intestine of animals exposed to the indicated bacteria for 24 h at 25 °C. Representative of 3 biological replicates, n = 15 - 25 animals per biological replicate. **A. H.** Area under the curve (AUC) quantitative analysis of P*t24b8.5::gfp* expression in the intestine, normalized to *E. coli* control. Animals fed on the indicated bacteria for 24 h at 25 °C. Three biological replicates, n = 15 – 25 animals per biological replicate. \*\*\*\**p* ≤ 0.0001; \**p* ≤ 0.05, ordinary one-way ANOVA, Dunnett’s multiple comparisons test, compared to *E. coli* controls.

Animals that were mono-associated with distinct CeMbio bacteria showed differential P*t24b8.5::gfp* expression. At one extreme, *Pantoea nemavictus* induced P*t24b8.5::gfp* throughout the intestine, for an average ∼2.5-fold compared with reference animals (**Fig. 2C, F, and H**). At the other extreme, *C. scophthalmum* repressed P*t24b8.5::gfp* about 2-fold (**Fig. 2D, G, and H**). The rest fell somewhere in between, with *L. amnigena, P. berkeleyensis,* inducing P*t24b8.5::gfp* about 2-fold (high induction), *E. hormaechei, A. guillouiae, S. molluscorum,* and *P. lurida* about 1.5-fold (modest induction), *S. multivorum* and *S. indicatrix* about 1-fold (no induction), and *O. vermis* slightly repressing (**Fig. 2F, G, and H**). These results suggested that the CeMbio bacteria may vary in their activation of the PMK-1/p38 MAPK pathway and hinted that it may be important for their interactions with *C. elegans*.

### PMK-1 mediates host defense against distinct microbiota members

To assess the role of PMK-1/p38 MAPK in interactions with CeMbio members, we performed survival assays on individual species, comparing *pmk-1-*deficient animals to wild type. First, we identified two informative times that enabled clear distinction of bacteria that promoted or impaired survival, in both backgrounds. As references, the median survival of wild type and *pmk-1(-)* animals exposed to nonpathogenic *E. coli* on NGM medium was about 7 d; starved animals (no bacteria) on TSA medium showed lifespan extension (median survival of 11 d); and *S. aureus-*infected animals showed decreased survival (medians of 2 and 1 d for wild type and *pmk-1(-),* respectively) (**Fig. 3A and B**). The differences among *S. multivorum*, *C. scophthalmum*, and *S. indicatrix* were best resolved at days 3 and 7, for both *C. elegans* genetic backgrounds (**Fig. 3A and B**). Thus, we chose these two time points for further study.

**Figure 3.**
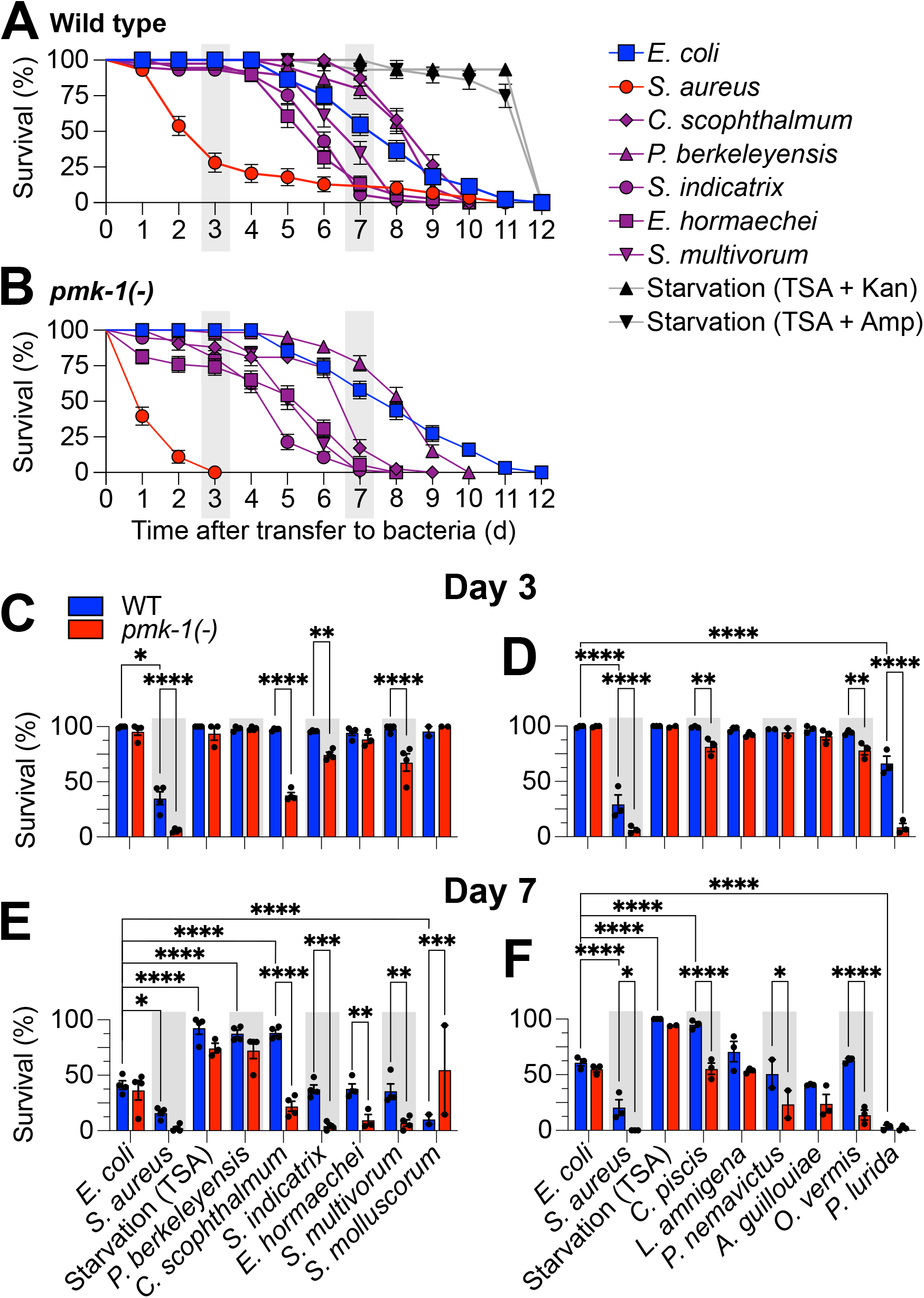
PMK-1/p38 MAPK is differentially required for *C. elegans* survival on distinct CeMbio bacteria. **A and B.** Survival of wild type (A) or *pmk-1(km25)* (B) animals after transfer to full lawns of CeMbio bacterial monoculture on TSA at 25 °C with the indicated antibiotics to prevent *E. coli* contamination. Data are representative of two biological replicates. Error bars are ± SEM. n = 50 - 100 animals per condition per replicate. **C – F.** Survival of wild type and *pmk-1(km25)* animals after transfer to full lawns of CeMbio bacterial monoculture on TSA at 25 °C on days 3 (C and D) and 7 (E and F). Trials were performed in two internally controlled batches as shown. Data points represent median survival from individual biological replicates; at least 2 biological replicates for each treatment. Error bars are ± SEM. n = 50 - 100 animals per condition per replicate. \*\*\*\**p* ≤ 0.0001, \*\*\**p* ≤ 0.001, \*\**p* ≤ 0.01, \**p* ≤ 0.05, ordinary two-way ANOVA, Dunnett’s multiple comparisons test.

We measured survival after 3 and 7 days of exposure to the individual species of the CeMbio set, benchmarking against animals associated with nonpathogenic *E. coli*. For wild type animals, *P. nemavictus, L. amnigena, E. hormaechei, A. guillouiae, S. multivorum, S. indicatrix,* and *O. vermis* supported lifespan that was indistinguishable from control (**Fig. 3E and F**). We categorized these as “Neutral”. *P. berkeleyensis, C. piscis, and C. scophthalmum* supported increased lifespan, while *P. lurida* and *S. molluscorum* shortened lifespan (**Fig. 3E and F**). We categorized these as “Probiotics” and “Pathogen”, respectively. Thus, we identified *S. molluscorum* and *P. lurida* as CeMbio species that in mono-association may exhibit virulence towards wild type *C. elegans.* Consistent with this interpretation, *pmk-1* deletion further decreased survival on *P. lurida* (**Fig. 3D, F**), supporting the notion that PMK-1/p38 MAPK is important for host defense against virulence expressed by *P. lurida*.

Among the Probiotics, *pmk-1* deletion had no effect on host survival with *P. berkeleyensis,* showing that PMK-1/p38 MAPK is dispensable for lifespan extension by that bacterium. In contrast, *pmk-1* deletion reduced survival with *C. scophthalmum* and *C. piscis,* suggesting that PMK-1/p38 MAPK is important for lifespan promotion by those two species (**Fig. 3E, F**). Remarkably, *pmk-1(-)* mutants fared worse than wild type on five of seven Neutrals, with the exceptions of *A. guillouiae* and *S. molluscorum* (**Fig. 3E, F**). Thus, PMK-1/p38 MAPK may play differential roles in maintaining or promoting lifespan (but not reducing it) when *C. elegans* is mono-associated with distinct members of its microbiota.

Taken together with the previous reporter expression data, these survival data defined four broad categories of CeMbio species (**Table 2**): Category 1, PMK-1 activated and required for defense (*P. nemavictus*, *P. lurida*, *C. piscis,* and *E. hormaechei*); Category 2, PMK-1 activated but not required for defense (*P. berkeleyensis*, *A. guillouiae*, *S. molluscorum* and *L. amnigena*); Category 3, PMK-1 not activated but required for defense (*S. multivorum* and *S. indicatrix*); and Category 4, PMK-1 repressed but required for survival (*O. vermis* and *C. scophthalmum*). These results demonstrate how activation of PMK-1/p38 MAPK signaling may not correlate well with its genetic requirement for protection against microbiota bacterial species.

### CeMbio bacteria differentially activate an ACh-WNT gene reporter in the intestinal epithelium

To assess ACh-WNT signaling, we used a previously characterized genetically-encoded P*clec-60::gfp* fluorescent reporter. This reporter is rapidly induced in the intestinal epithelium in a pathogen- and ACh-WNT-dependent manner (Irazoqui *et al*., 2010a). Reference animals on nonpathogenic *E. coli* showed low P*clec-60::gfp* expression and only in the posterior-most of intestinal ring, consistent with prior reports (**Fig. 4A, E**). Also as reported, P*clec-60::gfp* expression extended anteriorly in animals infected with *S. aureus*, serving as positive control (Irazoqui *et al*., 2010a; Labed *et al*., 2018) (**Fig. 4B, E**). However, in contrast to previous reports where P*clec-60::gfp* was not induced by shorter starvation regimens (Labed *et al*., 2018), by 24 h of starvation P*clec-60::gfp* expression did increase, providing an additional positive control **(Fig. 4H).**

**Figure 4.**
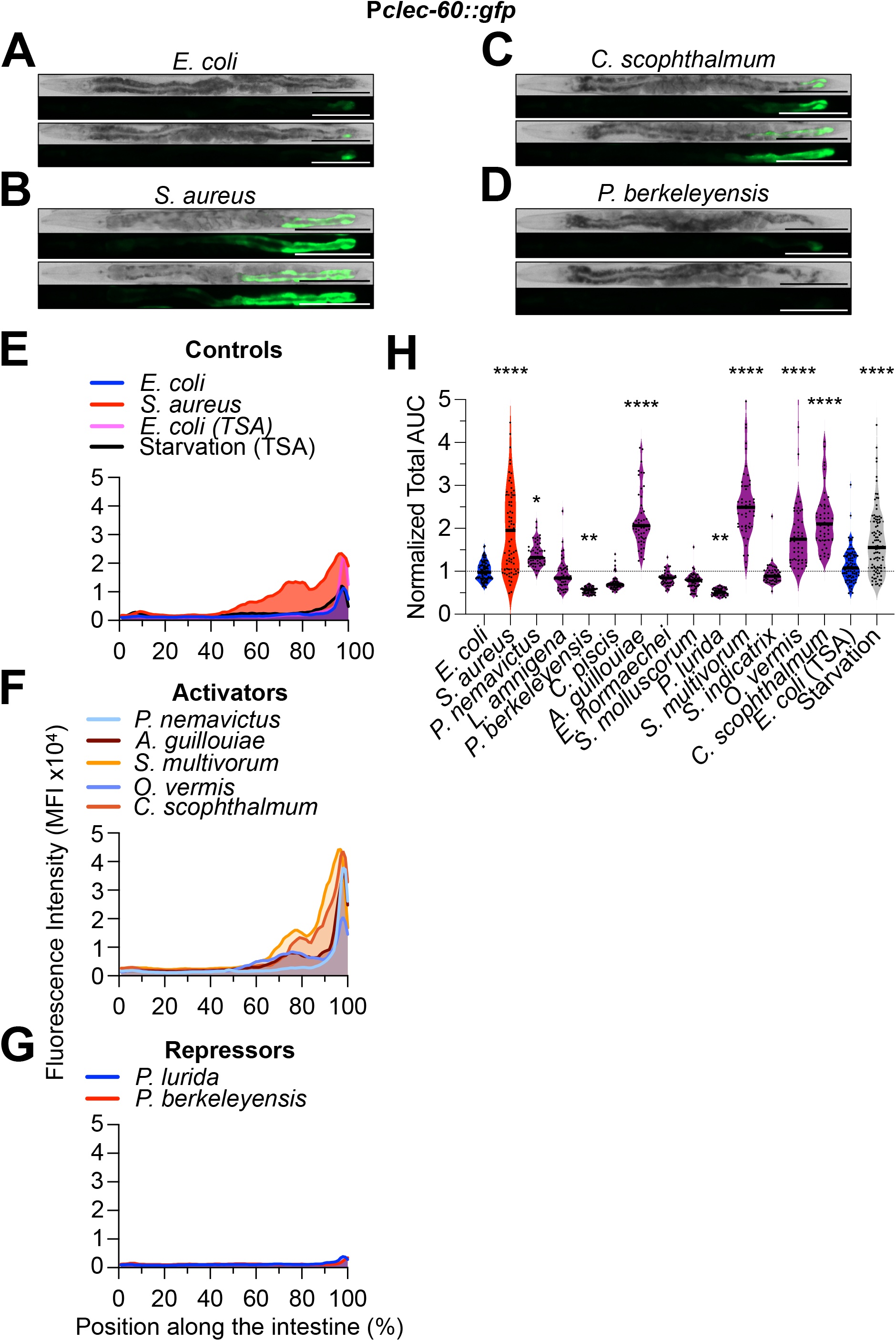
*C. elegans* microbiota bacteria differentially activate P*clec-60::gfp* expression along the intestinal epithelium. **A - D.** Representative brightfield and epifluorescence micrographs of P*clec-60::gfp* animals fed *E. coli*, *S. aureus*, *L. amnigena*, and *C. scophthalmum* for 24 h at 25 °C. Scale bars = 200 μm. **E - G.** P*clec-60::gfp* mean fluorescent intensity (MFI) along the intestine of animals exposed to the indicated bacteria for 24 h at 25 °C. Representative of 3 biological replicates, n = 15 - 25 animals per biological replicate. **H.** Area under the curve (AUC) quantitative analysis of P*clec-60::gfp* expression in the intestine, normalized to *E. coli* control. Animals fed on the indicated bacteria for 24 h at 25 °C. Three biological replicates, n = 15 – 25 animals per biological replicate. \*\*\*\**p* ≤ 0.0001, \*\*\**p* ≤ 0.001, \*\**p* ≤ 0.01, \**p* ≤ 0.05, ordinary one-way ANOVA, Dunnett’s multiple comparisons test, compared to *E. coli* controls.

As with P*t24b8.5::gfp,* P*clec-60::gfp* expression stratified the CeMbio bacteria in four broad categories. At one extreme, the “Activators” *A. guillouiae, S. multivorum,* and *C. scophthalmum* induced P*clec-60::gfp* expression anteriorly in the intestine, for an average of ≥ 2-fold compared to reference (**Fig. 4C, F, and H**). At the other extreme, the “Repressors” *P. berkeleyensis* and *P. lurida* repressed P*clec-60::gfp* expression about 1.5-fold (**Fig. 4D, G, and H**). The remaining bacteria fell in between. Aside from the “Non Inducers”, *O. vermis* and *P. nemavictus* (“Modest Inducers”) significantly but weakly induced P*clec-60::gfp* compared with controls (**Fig. 4F and H**). These results suggested that the ACh-WNT pathway may be differentially activated by distinct members of the CeMbio. However, because 24 h starvation also induced P*clec-60::gfp*, these results alone did not allow us to discriminate whether the inducer bacteria did so due to ACh-WNT pathway activation or due to poor nutrition. For this reason, and because not many bacteria activated *clec-60,* we chose to examine a third host defense pathway.

### CeMbio bacteria differentially activate HLH-30/TFEB in the intestinal epithelium

To examine the activation of HLH-30/TFEB, we used a previously characterized construct that tags HLH-30 with GFP (HLH-30::GFP) (Visvikis *et al*., 2014). In previous work we showed that HLH-30::GFP concentrates in cellular nuclei throughout the animal just 30 min after infection with *S. aureus* (Visvikis *et al*., 2014). Reference animals on nonpathogenic *E. coli* showed little HLH-30::GFP nuclear localization, both in the intestine and the rest of the body (**Fig. 6A**). *S. aureus* induced strong nuclear localization throughout the entire body, serving as positive control (**Fig. 6B**). Remarkably, all of the CeMbio bacteria activated HLH-30::GFP in the intestinal epithelium (**Fig. 6E**). This activation could also be systemic, as most bacteria activated HLH-30::GFP also in the head (**Fig. 6F**), with the exceptions of *S. molluscorum* and *S. multivorum* (**Fig. 6C, F**). Thus, HLH-30/TFEB activation is a hallmark of mono-association with the CeMbio bacteria, suggesting that HLH-30/TFEB plays a fundamental role in interactions between *C. elegans* and its microbiota.

**Figure 5.**
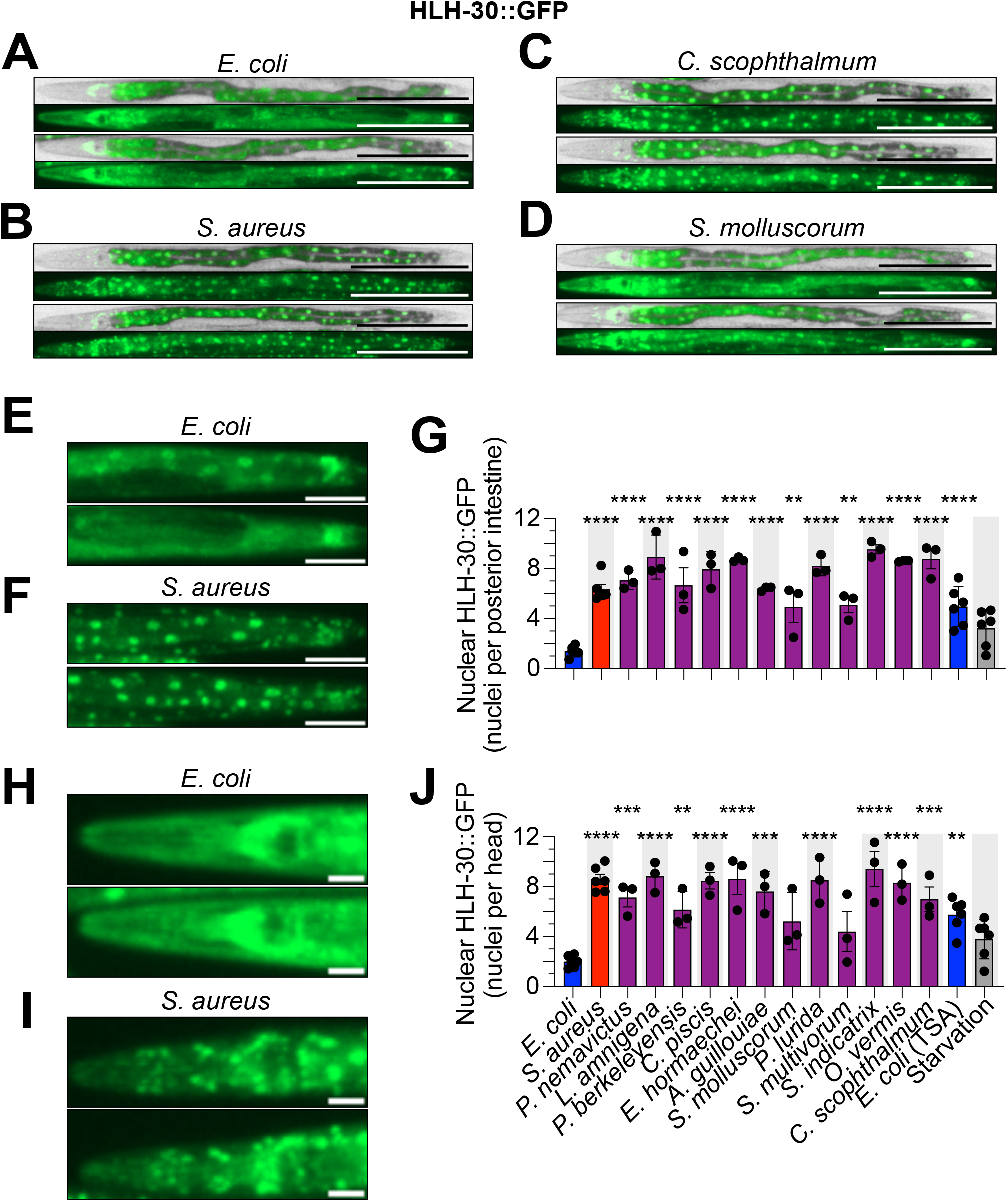
*C. elegans* microbiota bacteria differentially drive HLH-30::GFP nuclear localization. **A - D.** Representative brightfield and epifluorescence micrographs of P*hlh-30::hlh-30a::gfp* (HLH-30::GFP) animals fed *E. coli*, *S. aureus, S. molluscorum*, and *C. scophthalmum* for 30 min at 25 °C. Scale bars = 200 μm. E and F) Representative epifluorescence micrographs of HLH-30::GFP animals posterior intestine fed *E. coli* or *S. aureus* for 30 min at 25 °C. Scale bars = 50 μm H and I) Representative epifluorescence micrographs of HLH-30::GFP animals head fed *E. coli* or *S. aureus* for 30 min at 25 °C. Scale bars = 20 μm **G and H.** Quantitative analysis of HLH-30::GFP nuclear localization in the intestinal epithelium (G) or head (H) of animals fed the indicated bacteria for 30 min. Data are means ± SEM for three biological replicates; n = 15 - 25 animals per biological replicate. \*\*\*\**p* ≤ 0.0001, \*\*\**p* ≤ 0.001, \*\**p* ≤ 0.01, \**p* ≤ 0.05, ordinary one-way ANOVA, Dunnett’s multiple comparisons test, compared to *E. coli* controls.

**Figure 6.**
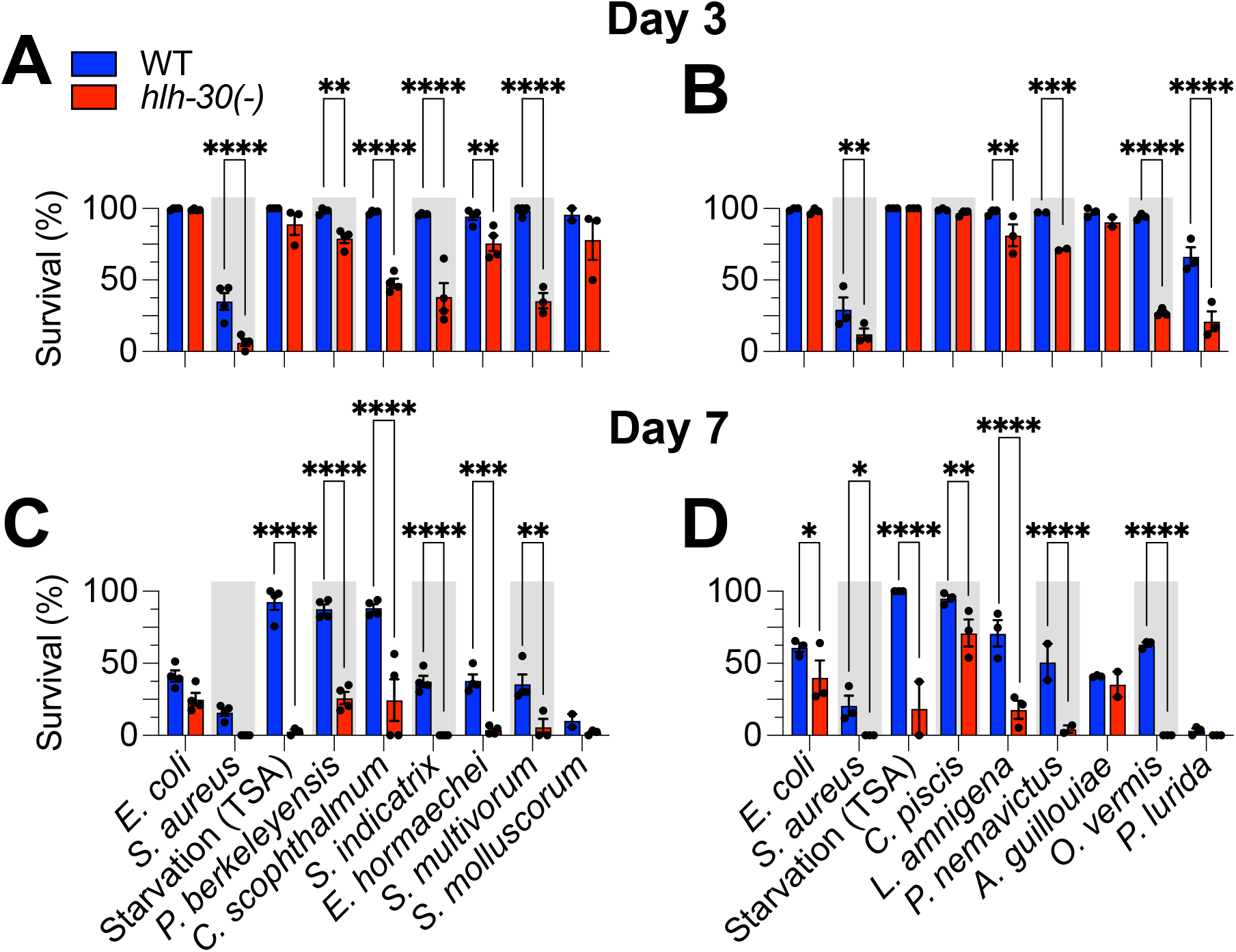
HLH-30 is differentially required for *C. elegans* survival on distinct CeMbio bacteria. **A – D.** Survival of wild type and *hlh-30(tm1978)* animals after transfer to full lawns of CeMbio bacterial monoculture on TSA at 25 °C on days 3 (A and B) and 7 (C and D). Trials were performed in two internally controlled batches as shown. Data points represent median survival from individual biological replicates; at least 2 biological replicates for each treatment. Error bars are ± SEM. n = 50 - 100 animals per condition per replicate. \*\*\*\**p* ≤ 0.0001, \*\*\**p* ≤ 0.001, \*\**p* ≤ 0.01, \**p* ≤ 0.05, ordinary two-way ANOVA, Dunnett’s multiple comparisons test.

### HLH-30/TFEB promotes host survival in interactions with distinct microbiota members

Following the same approach as for PMK-1/p38 MAPK, we used *hlh-30(-)* mutants to assess the role of HLH-30/TFEB in interactions with members of its CeMbio microbiota. The *E. coli-*associated references for both genotypes were indistinguishable at day 3 but could be resolved by day 7, as expected, given the known aging defect exhibited by *hlh-30*(-) animals (Visvikis *et al*., 2014). The starved *hlh-30(-)* controls showed a dramatic loss of viability at day 7, consistent with previous reports (Settembre *et al*., 2013).

At day 3, *hlh-30(-)* mutants fared equally well with *S. molluscorum, C. piscis* and *A. guillouiae,* as they did with *E. coli* (**Fig. 6A and C**). In contrast, the rest of the CeMbio bacteria reduced the survival of *hlh-30(-)* mutants relative to wild type at day 3, suggesting that HLH-30/TFEB is essential for host defense against these bacteria (**Fig. 6A and C**). At day 7, the relationships remained essentially the same, except for *C. piscis,* which caused earlier death of *hlh-30(-)* mutants compared to wild type (**Fig. 6B and D**). Thus, all the bacteria induced HLH-30/TFEB nuclear import and showed virulence towards *hlh-30-*deficient animals, except for *S. molluscorum* and *A. guillouiae,* which did not show virulence.

## *Acinetobacter guillouiae* promotes survival of infection with natural pathogen *Pseudomonas lurida*

*A. guillouiae* activated all three pathways and did not exhibit virulence towards *pmk-1(-)* or *hlh-30(-)* animals, suggesting that it may possess probiotic properties. In contrast, *P. lurida* moderately induced PMK-1/p38 MAPK, strongly activated HLH-30/TFEB, and exhibited virulence towards wild type, *pmk-1(-),* and *hlh-30(-)* animals, suggesting that *P. lurida* possesses the hallmarks of an overt (and natural) pathogen of *C. elegans*.

Because PMK-1/p38 MAPK and HLH-30/TFEB are activated by *P. lurida* and both pathways are required for defense against its virulence, we tested if their prior activation by *A. guillouiae* may be protective (**Fig. 7A**). A relatively short prior exposure of wild type animals to *A. guillouiae* significantly enhanced survival of *P. lurida* infection compared to *E. coli-*exposed controls (median survival ∼5 days v. ∼3 days, **Fig. 7B**), confirming that *A. guillouiae* protects *C. elegans* from *P. lurida* virulence.

**Figure 7.**
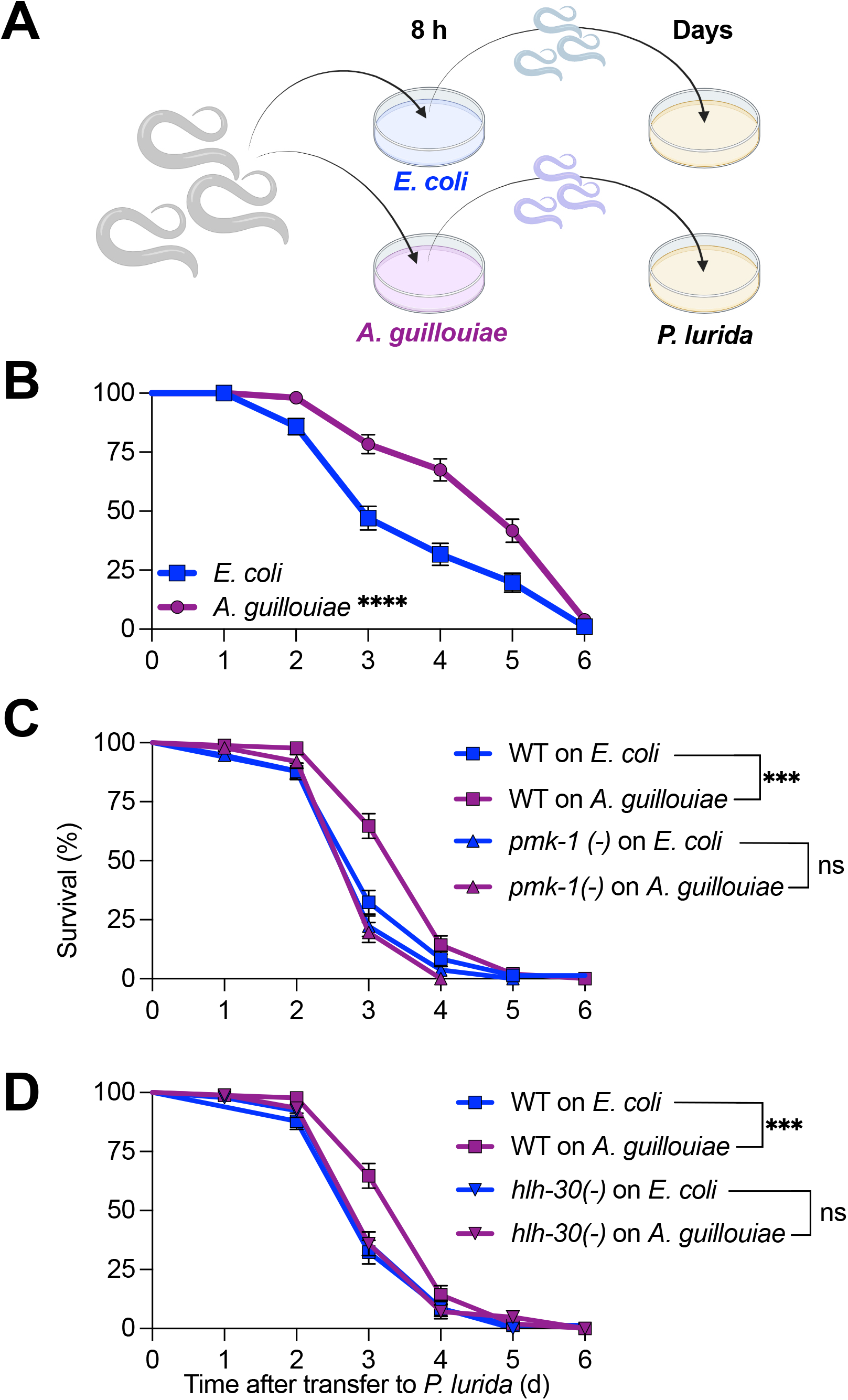
*A. guillouiae* exerts immune-mediated host protection against *P. lurida* infection. **A.** Schematic of the experimental approach. Animals were transferred from *E. coli* to *A. guillouiae* on TSA for 8 h at 25 °C. Subsequently, they were transferred to full lawns of *P. lurida* on TSA at 25 °C and scored for survival. In parallel, controls animals were transferred to *E. coli* instead of *A. guillouiae* and treated identically. **B.** Survival of *P. lurida* infection of wild type animals pre-exposed to *E. coli* or *A. guillouiae* for 8 h. Representative of 3 biological replicates, n = 50-100 animals. Error bars are ± SEM from 3 technical replicates. \*\*\*\**p* ≤ 0.0001, \*\*\**p* ≤ 0.001, \*\**p* ≤ 0.01, \**p* ≤ 0.05, Kaplan-Meier Log-rank (Mantel-Cox) test. **C.** Survival of *P. lurida* infection of pre-exposed wild type and *pmk-1(-)* animals. Representative of 3 biological replicates, n = 50-100 animals. Error bars are ± SEM from 3 technical replicates. \*\*\*\**p* ≤ 0.0001, \*\*\**p* ≤ 0.001, \*\**p* ≤ 0.01, \**p* ≤ 0.05, Kaplan-Meier Log-rank (Mantel-Cox) test. **D.** Survival of *P. lurida* infection of pre-exposed wild type and *hlh-30(-)* animals. Representative of 3 biological replicates, n = 50-100 animals. Error bars are SEM from 3 technical replicates. \*\*\*\**p* ≤ 0.0001, \*\*\**p* ≤ 0.001, \*\**p* ≤ 0.01, \**p* ≤ 0.05, Kaplan-Meier Log-rank (Mantel-Cox) test.

*A priori, A. guillouiae* could protect *C. elegans* directly, by displacing *P. lurida* from the intestinal lumen (niche competition model) or indirectly, by inducing host defense (third-party competition model). To discriminate between these scenarios, we performed *A. guillouiae* protection assays with *pmk-1(-)* and *hlh-30(-)* mutants. Remarkably, deletion of either *pmk-1* (**Fig. 7C**) or *hlh-30* **(Fig. 7D**) completely suppressed the protection conferred by *A. guillouiae.* These results confirmed that protection conferred by *A. guillouiae* against *P. lurida* requires, and is likely mediated by, both the PMK-1/p38 MAPK and the HLH-30/TFEB pathways.

Recall that “nonpathogenic” *E. coli* activated PMK-1/p38 MAPK and HLH-30/TFEB on TSA medium (**Fig. 2H and 6G).** Prior exposure to *E. coli* on TSA medium also protected against *P. lurida* **(Sup Fig. 2A)***. hlh-30* deletion animals showed protection towards *P. lurida* conferred by *E. coli* (**Sup Fig. 2B**). Thus, in this case HLH-30/TFEB pathway was not required for the probiotic effect of *E. coli*.

## Discussion

We report here a phenotypic description of the reference *C. elegans* microbiota, known as CeMbio, including the morphological and growth characteristics of its members in vegetable-derived rich culture media in presence or absence of antibiotics, their ability to induce any of three important host defense pathways, and their ability to cause disease in animals that lack them. To our knowledge, this is the first systematic examination of these properties for every CeMbio member, at least under conditions that more closely resemble the natural niche of *C. elegans* than the standard nematode growth medium that is widely used in the laboratory.

The first key insight to emerge from these studies was that the ability of each bacterial species to induce reporters for a given host defense pathway did not correlate well with its virulence in wild type animals nor the requirement of that pathway for host defense against the bacteria (**Table 2**, **Fig. 8**). This is somewhat surprising, because a simple assumption might be that each host defense pathway evolved to detect and defend against bacteria that are virulent. We did find examples of bacteria that induced a reporter and were more virulent against animals that lacked the corresponding pathway, including some that showed “cryptic” virulence, or virulence that was masked by the (appropriate) host response. However, this was not the rule. What might be the evolutionary advantage to the host, or the inducing bacteria, to activate a host response if it does not protect the host against that infection? One possibility might be that the host response may protect the host against a second bacterial pathogen, enhancing the persistence and availability of the intestinal niche for the inducing bacteria.

**Figure 8.**
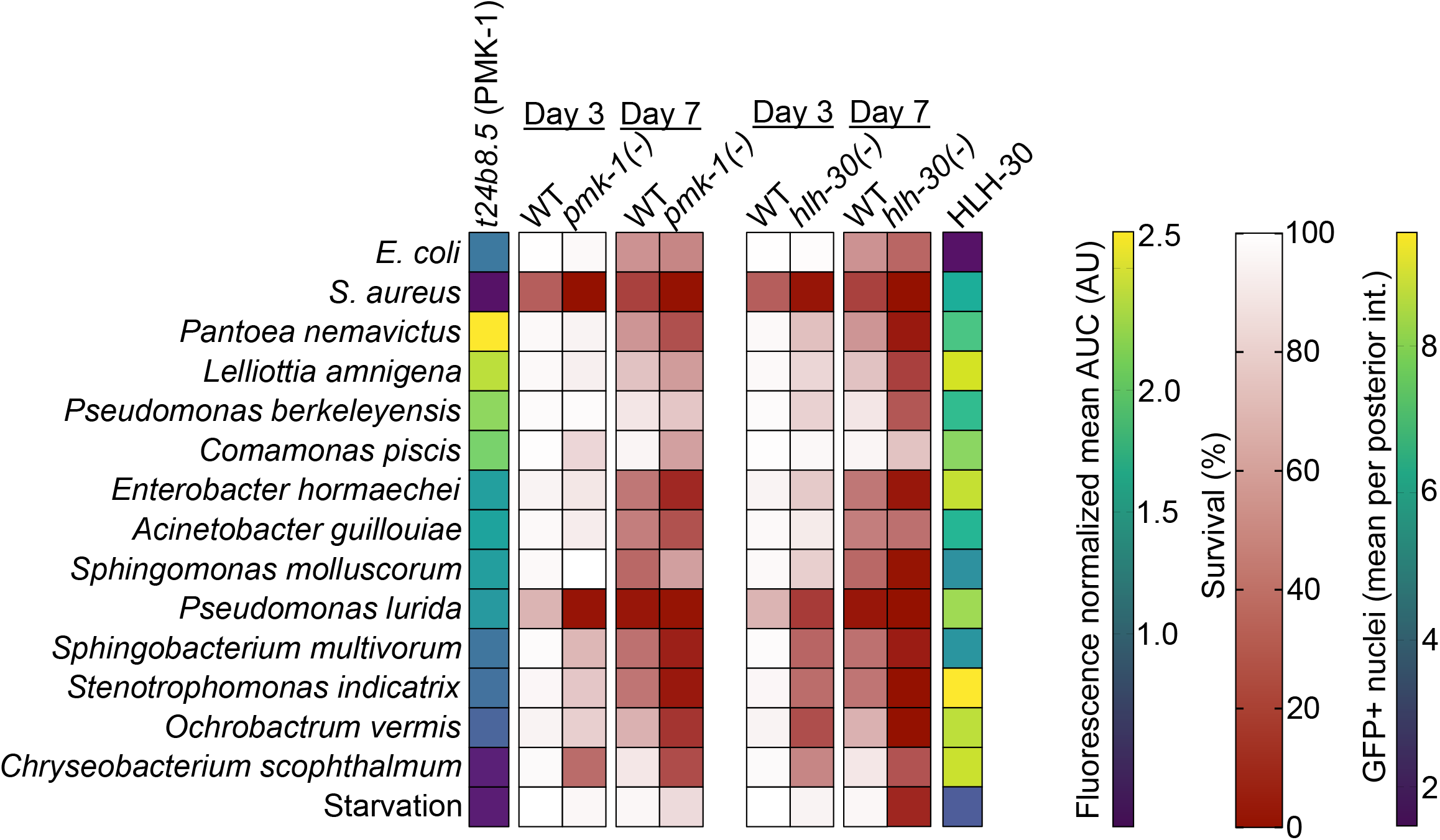
Summary of pathway analysis data. Reporter activity and survival at days 3 and 7 of animals exposed to the indicated CeMbio bacteria. Represented data are from Fig. 2 - 6. In the survival heat maps, the WT data are represented twice to facilitate comparison with the *pmk-1(-)* and *hlh-30(-)* mutants.

A second key insight was that many CeMbio bacteria did not affect the survival of the host unless host defense was compromised (**Fig. 8**). In these examples, the bacteria would not be characterized as frank pathogens because they do not impair the survival of wild type animals compared to controls. Instead, they would fall under a category known as “opportunistic” pathogens, revealing that they do indeed possess pathogenic potential that is masked in immunocompetent animals, what we call ‘cryptic’ virulence (Casadevall and Pirofski, 2003; Casadevall and Pirofski, 2015). On the other hand, over-induction of a host response pathway could conceivably lead to disease, or improved health after removal of the pathway.

The third key insight was the host protection against *P. lurida,* the strongest frank pathogen in our dataset, by *A. guillouiae* (**Fig. 7B**). Both bacteria have been previously shown to persist in the *C. elegans* intestine (Pees *et al*., 2021). Consistent with our finding that *P. lurida* exhibits virulence against *C. elegans,* the nematodes avoided *P. lurida* in food choice assays (Petersen *et al*., 2021). However, under certain conditions *P. lurida* may also provide protection against infection, at least against pathogenic fungi (Dirksen *et al*., 2016). *A priori,* protection against *P. lurida* pathogenesis by *A. guillouiae* could be mediated by colonization resistance (Sorbara and Pamer, 2019) or by immune-mediated protection (Kamada *et al*., 2013). Our data show that disruption of either the PMK-1/p38 MAPK or the HLH-30/TFEB pathway abrogates such protection, supporting an immune-mediated antagonistic relationship – a ‘Third-Party’ model of microbe-microbe interaction. This result suggests that the order of colonization by these microbiota members may be a key determinant of host health under natural conditions. Further investigation is warranted to define the extent to which direct niche competition may play a role as well. Moreover, it will be important to scale up this type of investigation into all potential microbe-microbe interaction pairs and, ultimately, entire communities *in vivo*.

Finally, our results clearly showed that HLH-30/TFEB becomes activated by ten out of twelve members of the CeMbio community. This observation suggests that HLH-30/TFEB is a key and broad element of microbiota-host interactions. It also suggests that there are commonalities among the ten activating species, which result in HLH-30/TFEB activation. It will be important to understand if these ten species are sensed by a common mechanism, or by distinct mechanisms that converge on HLH-30/TFEB. Future research should address the physiological consequences of HLH-30/TFEB activation on the host, on the bacteria, and on the community.

## Acknowledgments

Thanks to the members of the IMP graduate program and of the Department of Microbiology and Physiological Systems for helpful discussions. The Sanderson Center for Optical Experimentation of the University of Massachusetts Chan Medical School provided microscopy support. Special gratitude to Dr. Amanda Wollenberg for initial work with microbiota isolates, and to Amy Parker, Annette Bohigian, Richard Fish, Tracey Rae, Marie Berardi, and Dhruti Desai for exceptional administrative support. Research reported in this publication was supported by the National Institute of General Medical Sciences of the National Institutes of Health under award numbers R01GM101056 and R35GM149284 (JEI), by the National Institute of Allergy and Infectious Diseases of the National Institutes of Health under award numbers T32AI095213 (XG), T32AI095213 (XG), and R21AI169842 (JEI), and by the Dr. Marcellette G. Williams Memorial Fund (JEI). Some strains were provided by the CGC, which is funded by NIH Office of Research Infrastructure Programs (P40OD010440). The content is solely the responsibility of the authors and does not necessarily represent the official views of the National Institutes of Health.

## Experimental Procedures

### Strains, media, and culture conditions

*C. elegans* strains were maintained on nematode-growth media (NGM) plates seeded with *E. coli* OP50 at 15 – 20°C, according to standard procedures (Powell and Ausubel, 2008). See **Table 4** for strain genotype details. The *C. elegans* microbiota kit was purchased through the Caenorhabditis Genetics Center (CGC).

CeMbio cultures were grown under aerobic conditions at 25 °C with 170-200 RPM shaking in TSB (Sigma-Aldrich T8907) containing 50 µg ml^-1^ ampicillin for 20-24 h. 10 μL of the CeMbio culture was then spread on TSA (BD 236950) containing 50 µg ml^-1^ ampicillin plates for 20-24 h at 25 °C. TSA containing 50 µg ml^-1^ ampicillin plates were made within 10 days of use for experiments. Tryptic soy agar (TSA), as a plant hydrolysate, is a closer approximation to the natural environment from which they were isolated than other more traditional media (e.g., NGM or Luria-Bertani) (Stiernagle, 2006).

*S. aureus* SH1000 was grown overnight in TSB containing 50 µg ml^-1^ kanamycin, 37 °C at 200 RPM. 10 μL of overnight cultures was spread on the surface of 35 mm TSA containing 10 µg ml^-1^ kanamycin and incubated 5–6 h at 37 °C. Animals were transferred to each of three replicate infection plates. Survival was quantified as described (Powell and Ausubel, 2008).

For *E. coli* OP50 grown on TSA plates, *E. coli* was grown overnight in LB (MP Biomedicals 113002032) containing 100 µg ml^-1^ streptomycin, 37 °C at 170-200 RPM. 10 µL of overnight culture was spread on the surface of 35 mm TSA containing 50 µg ml^-1^ ampicillin, and incubated 25 °C, similarly to CeMbio. Control NGM plates were seeded with 200 µL of overnight culture and incubated at 25 °C.

### Survival

For *hlh-30* and *pmk-1* mutant animals: Animals were staged by transferring egg laying adults to *E. coli* NGM plates and allowed to lay eggs for 2-3 hours. Egg laying adults were then removed. Progeny were placed at 15 °C overnight and then shifted to 20 °C for 2 days, reaching L4/Young adult (YA) stage. YA animals were washed 10x in 10-14mL 1xM9 buffer and transferred to UV-arrested E. coli (UV-arrested to minimize carry over of E. coli in subsequent steps). YA animals on UV-*E. coli* were grown at 25C overnight (*spe-9*(hc88) I; *rrf-3*(b26) II mutants are fertilization defective at 25°C. The adult animals are then transferred onto full lawn plates of *S. aureus*, *E. coli*, starvation, and CeMbio plates, 3-2 technical replicates per genotype.

### A. guillouiae protection

L4 animals were placed at 25 °C for 20-24 h to reach YA stage. ∼100 YA animals were then placed on *E. coli* NGM/TSA plates or A. *guillouiae* for 8 h at 25 °C. After 8 h, ∼30 animals were transferred on *P. lurida* plates in triplicate, placed at 25 °C, and scored every day.

### Image acquisition and analysis

Images were taken using a Lionheart FX automatic microscope (BioTek Instruments) using 4x objective. 10-25 YA animals were placed onto bacterial plates and imaged after 30 min for HLH-30::GFP, 4 h for P*fmo-2::gfp*, 24 h for P*t24b8.5::gfp* and P*clec-60::gfp* at 25 °C. Fluorescence microscopy experiments were repeated total of 3 independent times. Animals were anesthetized using 100 mM NaN_3_ (Sigma-Aldrich S2002) on a 4% agar pad for imaging. All images were captured at the same exposure and intensity in a biological replicate. Greyscale images were used for image analysis. FIJI was used for image analysis. A region of interest (ROI) was drawn using segmented line tool along the intestine or whole animal, from anterior to posterior.

For P*t24b8.5::gfp*, and P*clec-60::gfp*, mean fluorescent intensity (MFI) of the ROI, anterior to posterior intestine, was determined by using the Analyze>Multi Plot function. Pixel length was transformed to percent length in excel, each percent length value has a single value. All percent length-MFI values were rounded up to the nearest whole percent length number in excel. Area under the curve was calculated using GraphPad PRISM9 and normalized to *E. coli* control.

For HLH-30::GFP animals, an ROI of the head (from mouth to pharyngeal bulb) and the anterior half of intestine was acquired. A threshold was applied to images by converting RAW grayscale image to 8bit. The threshold was determined by the most HLH-30::GFP nuclei on *S. aureus* and least HLH-30::GFP nuclei on *E. coli* treated animals. The threshold is the same within a biological replicate. ROI’s were selected using ROI Manager>”OR (Combine)” and analyzed using Analyze Particles. Analyze Particles set values are Size=1-20, Circularity=0-1.

For single colony pictures, bacteria were streaked out on TSA + 50 µg ml^-1^ Ampicillin plates and placed at 25 °C until colonies were seen, 24 – 48 h. Pictures were taken once single colonies were visible using iPhone 11 camera through dissecting microscope eyepieces on 3.5x zoom.

For antibiotic resistance, ampicillin, erythromycin, kanamycin, and streptomycin were used. The antibiotics was added once autoclaved TSA media cooled and then added to single well plates. CeMbio was grown in TSB at 25C for 24 h, and 2 µL of culture was pipetted onto the TSA + antibiotic plates. CeMbio were placed at 25 °C for 24 h and picture was taken using iPhone 11 camera.

### Statistics

Statistics analyses were performed in GraphPad PRISM9. For comparison of survival curves, a Log-rank (Mantel-Cox) test was performed. For % survival on CeMbio at days 3 and 7, an ordinary two-way ANOVA, Dunnett’s multiple comparisons test was performed. For fluorescence analyses, an ordinary one-way ANOVA, Dunnett’s multiple comparisons test, compared to *E. coli* controls was performed. A p-value ≤ 0.05 was considered significantly different from control.

## Figure legends

**Supplemental Figure 1. Antibiotic resistance of CeMbio bacteria.**

**A.** Growth on TSA with indicated antibiotics of 2 μl overnight aliquots for 24 h at 25 °C.

**Supplemental Figure 2. *E. coli* on TSA protects against *P. lurida*.**

**A.** Survival of *P. lurida* infection of wild type animals pre-exposed to *E. coli* on NGM or TSA for 8 h. Representative of 2 biological replicates, n = 50-100 animals. Error bars are ± SEM from 3 technical replicates. \*\**p* ≤ 0.01, \**p* ≤ 0.05, Kaplan-Meier Log-rank (Mantel-Cox) test.

**B.** Survival of *P. lurida* infection of *hlh-30(-)* deficient animals pre-exposed to *E. coli* on NGM or TSA for 8 h. Representative of 2 biological replicates, n = 50-100 animals. Error bars are ± SEM from 3 technical replicates. \**p* ≤ 0.05, Kaplan-Meier Log-rank (Mantel-Cox) test.

**Table 1. Antibiotic susceptibility of the 12 CeMbio isolates**

**Table 2. Categorization of CeMbio isolates according to reporter expression and effect of PMK-1 deletion on survival**

**Table 3. Categorization of CeMbio isolates according to reporter expression and effect of HLH-30 deletion on survival**

**Table 4. *C. elegans* strains used**

