## Supplementary figures and images for "Distinct members of the *C. elegans* CeMbio reference microbiota exert cryptic virulence and infection protection"

### Supplemental Figures

#### Gonzalez et al Sup Figure 1

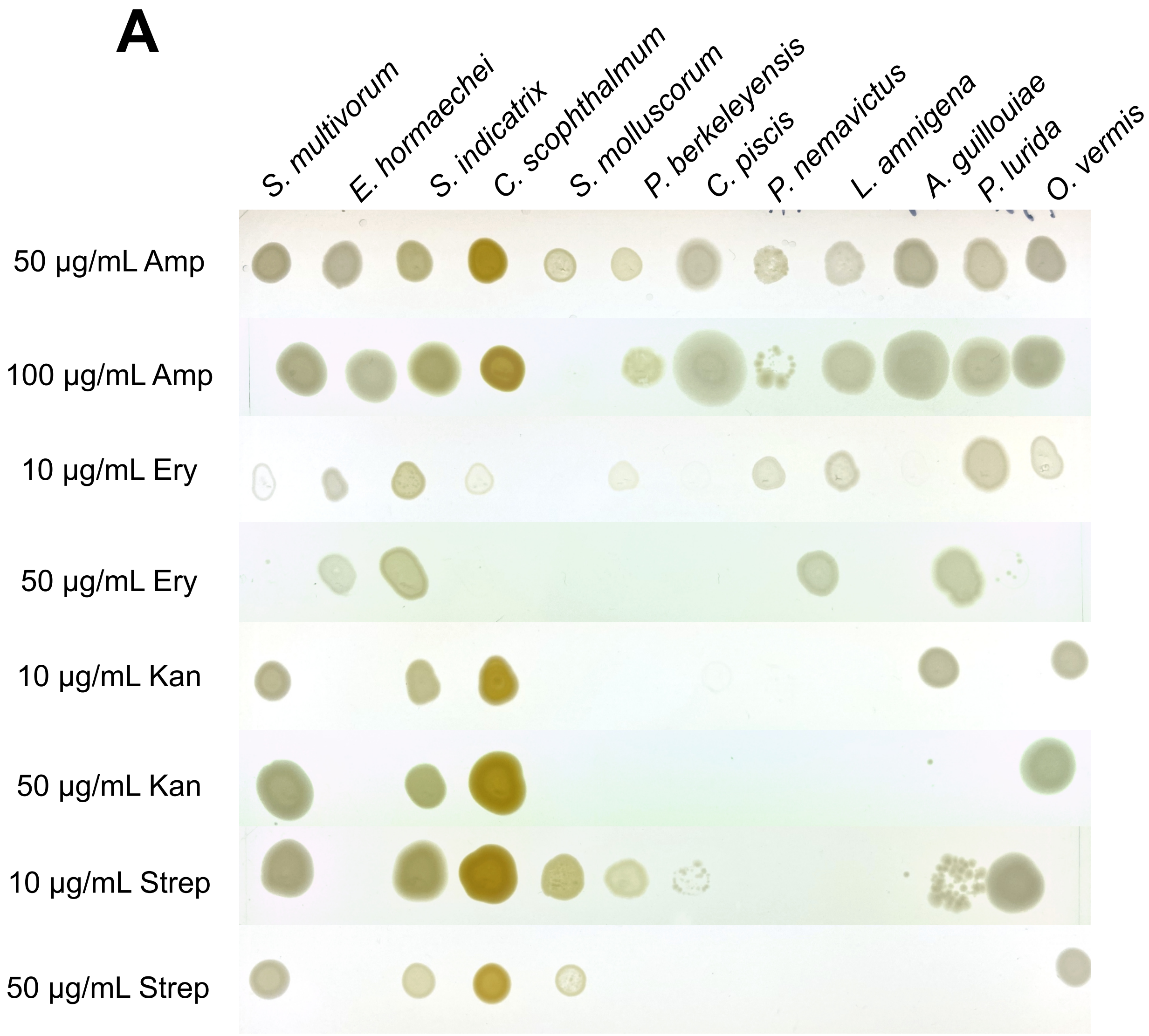

#### Gonzalez et al Sup Figure 2

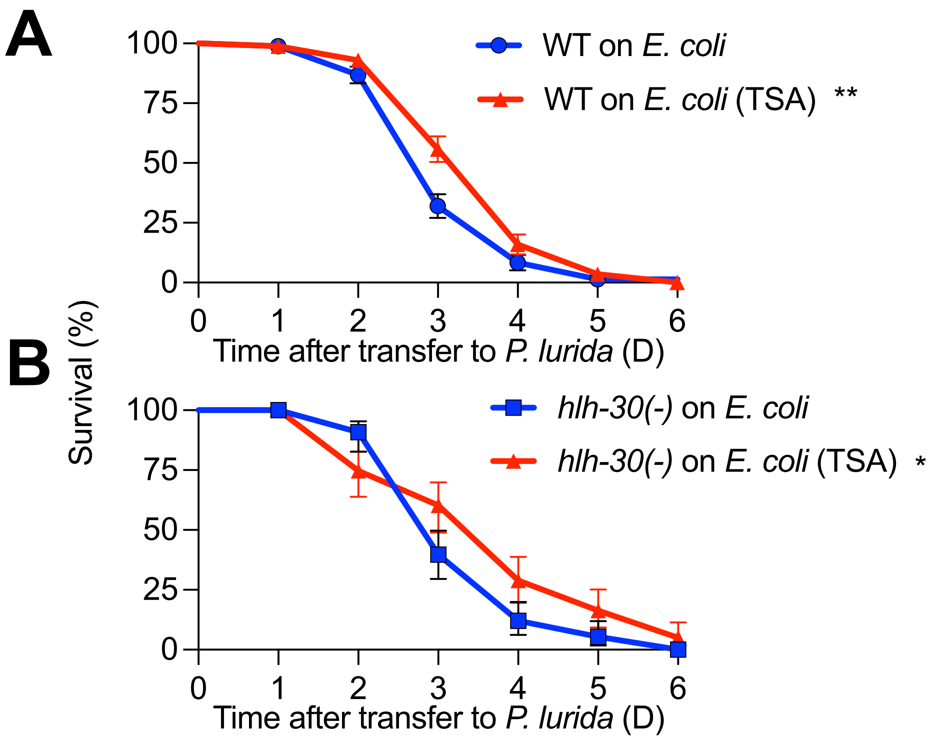
